# The cage effect of electron beam irradiation damage in cryo-electron microscopy

**DOI:** 10.1101/2024.03.20.585930

**Authors:** Yi Li, Dong-Dong Kang, Jia-Yu Dai, Lin-Wang Wang

## Abstract

Electron beam irradiation can cause damage to biological and organic samples, as determined via transmission electron microscopy (TEM). Cryo-electron microscopy (cryo-EM) significantly reduces such damage by quickly freezing the environmental water around organic molecules. However, there are multiple hypotheses about the mechanism of cryo-protection in cryo-EM. A lower temperature can cause less molecular dissociation in the first stage, or frozen water can have a “cage” effect by preventing the dissociated fragments from flying away. In this work, we used real-time time-dependent density functional theory (rt-TDDFT-MD) molecular dynamic simulations to study the related dynamics. We used our newly developed natural orbital branching (NOB) algorithm to describe the molecular dissociation process after the molecule is ionized. We found that despite the difference in surrounding water molecules at different temperatures, the initial dissociation process is similar. On the other hand, the dissociated fragments will fly away at room temperature, while they will remain in the same cage when frozen water is used. Our results provide direct support for the cage effect mechanism.

## Introduction

Since its development in the 1980s, cryo-electron microscopy (cryo-EM) has become one of the most useful tools in the field of biology^1^. This approach allows direct observation of biological structures such as proteins^2^. Recently, cryo-EM has also been used in materials research, especially for observing soft organic materials^3–5^. Compared with conventional transmission electron microscopy (TEM), where specimen damage can be significant, cryo-EM, developed by Dubochet and colleagues^6,7^ freezes specimens using liquid nitrogen and has been found to significantly reduce specimen damage during microscopic observation^6–9^. However, there has been controversy about the mechanisms underlying this damage reduction. One theory suggests that during the initial stage of ionization, low temperature can reduce bond breaking, resulting in a decreased number of fragments being produced^10,11^. An alternative theory proposes that the initial ionization and bond breaking might be the same; however, frozen surrounding water can form a cage, which prevents the dissociated fragments from flying away; thus, they can self-heal later by reacting back to their original molecule^9,12,13^. Unfortunately, direct experimental evidence for distinguishing these two mechanisms is difficult, which has made this a controversial issue for a long time.

There are several methods for inducing material damage during electron beam-induced radiolysis. For bulk solid materials, one way is direct elastic knock-off of the nucleus from its crystal position^14^. However, for small organic molecules, a more efficient method is ionization of the molecule by knocking out a deep electron from the molecule^15–17^. The initial ionization process is followed by hot carrier cooling, which can empty the bonding electronic state and transfer the hot carrier electron energy to the nucleus, hence causing bond breakage^18,19^. Here, we study the molecular ionization-induced radiolysis process. The initial ionization is caused by the Coulomb interaction between the valence electron in the molecule and the fast-passing electron from TEM (which can have an energy ranging from a few thousand eV to millions of eV). This process is rather like a light absorption process, with the same oscillator strength^20–23^. A bound valence electron can absorb energy from a passing beam electron, be pumped out of the molecule, and be left with a molecule with a deep hole. This initial direct ionization process does not involve phonons and thus should have a rather insignificant dependence on the temperature and the surrounding environment; therefore, we can assume that they are the same for the room temperature and frozen temperature cases. Consequently, our simulations will start from this initial ionized molecule stage.

Sanche and coworkers^24–26^ reported that bond breaking begins with electron relaxation and subsequent energy distribution to vibrational modes. However, these studies are based on the gas phase and thus do not reveal the effect of the aqueous environment. Previous studies have also shown that samples in aqueous solutions can undergo less damage than isolated molecules^27,28^. Therefore, it is crucial to conduct investigations with aqueous environments under different conditions. In our simulations, we chose an organic molecule embedded in a box of water molecules. Ab initio molecular dynamics will be used to determine the equilibrium atomic configurations. At room temperature, the water molecules are in the liquid phase, while at liquid nitrogen temperature, they freeze into an ice-like disordered state. In the following study, we will use the ethylene carbonate (EC) molecule as our test molecule.

Having set up the initial simulation system with their atomic configuration and electron ionization, we then used rt-TDDFT-MD to simulate the hot carrier relaxation and molecular bond breaking process^29–32^. This chemical reaction induced by excited electron state relaxation is often simulated by quantum chemical methods, such as time-dependent configuration interaction single calculations and coupled-cluster methods, without ionic dynamics^33–35^. However, such quantum chemical calculations can only address small molecules; thus, dealing with environmental water molecules is difficult. In contrast, the modern rt-TDDFT-MD method allows us to simulate a few hundred atoms and a few picoseconds^36,37^. However, a straightforward rt-TDDFT-MD simulation follows the mean-field Ehrenfest dynamics^38–40^. These dynamics lack two crucial aspects for studying hot carrier relaxation processes^41–46^. The first is the lack of detailed balance. As a result, a hot carrier cannot be efficiently cooled down. The second aspect is the stochastic nature of the dynamics. As a result, deterministic processes are difficult to use to statistically determine the probabilities of different reaction paths. Here, we used the newly developed natural orbital branching (NOB) formalism of the rt-TDDFT-MD, which is implemented in PWmat software^47–49^. The NOB takes into account the dephasing time due to electron and phonon wavefunction coupling and then uses the wavefunction collapsing approximation to describe the dephasing process. One unique feature of NOB is that the system can collapse into a set of natural orbitals of the density matrix instead of the usual eigenenergy orbital set. The wavefunction collapsing happens stochastically, and the requirement to compensate for the energy by the reaction degree of freedom’s nuclear kinetic energy restores the detailed balance. Overall, NOB is a classical Newton dynamic treatment for nuclear movements, and the quantum mechanical time-dependent Schrodinger equation is used for electron wavefunctions. It uses the wavefunction collapsing approach to approximate dephasing processes (which fundamentally arise from the quantum mechanical treatment of nuclear movements). It’s worth noting that none of the broken-down fragments is single H, thus the quantum effect of H movement will not significantly influence the fragment movement, and we have not included the quantum effect in either the NOB or AIMD simulations.

## Results

To include the aqueous environment while considering computational cost, we constructed a 14*14*14 A^3^ box that included one organic EC molecule and 88 water molecules. This structure is displayed in Fig. 1(a), where water molecules cluster around the EC molecule, resulting in a density of approximately 0.94g/cm^3^, corresponding to the density of low-density amorphous ice (LDA) in frozen samples, as measured in the experiment^50,51^. To achieve a reasonably disordered system, we initially heated a hexagonal ice-phase system to 1000 K. Notably, no bond breaking of water molecules occurred during this process. Subsequently, we annealed the disordered structures separately to 100 K (frozen conditions, near liquid nitrogen temperatures) and 300 K (room temperature conditions), which is more in line with the RDF results obtained from simulations^50,51^, as shown in Fig. 1(b) and Fig. S1 in the Supplemental Material. We then performed molecular dynamics relaxation calculations at the respective temperatures and DFT calculations for the simulated system; the energy level information, as well as the information of the main few electronic wavefunctions, is shown in Fig. 1(c)(d). Only after equilibrium was established did we begin the rt-TDDFT-MD calculations in microcanonical ensembles^52^.

**Fig. 1.**
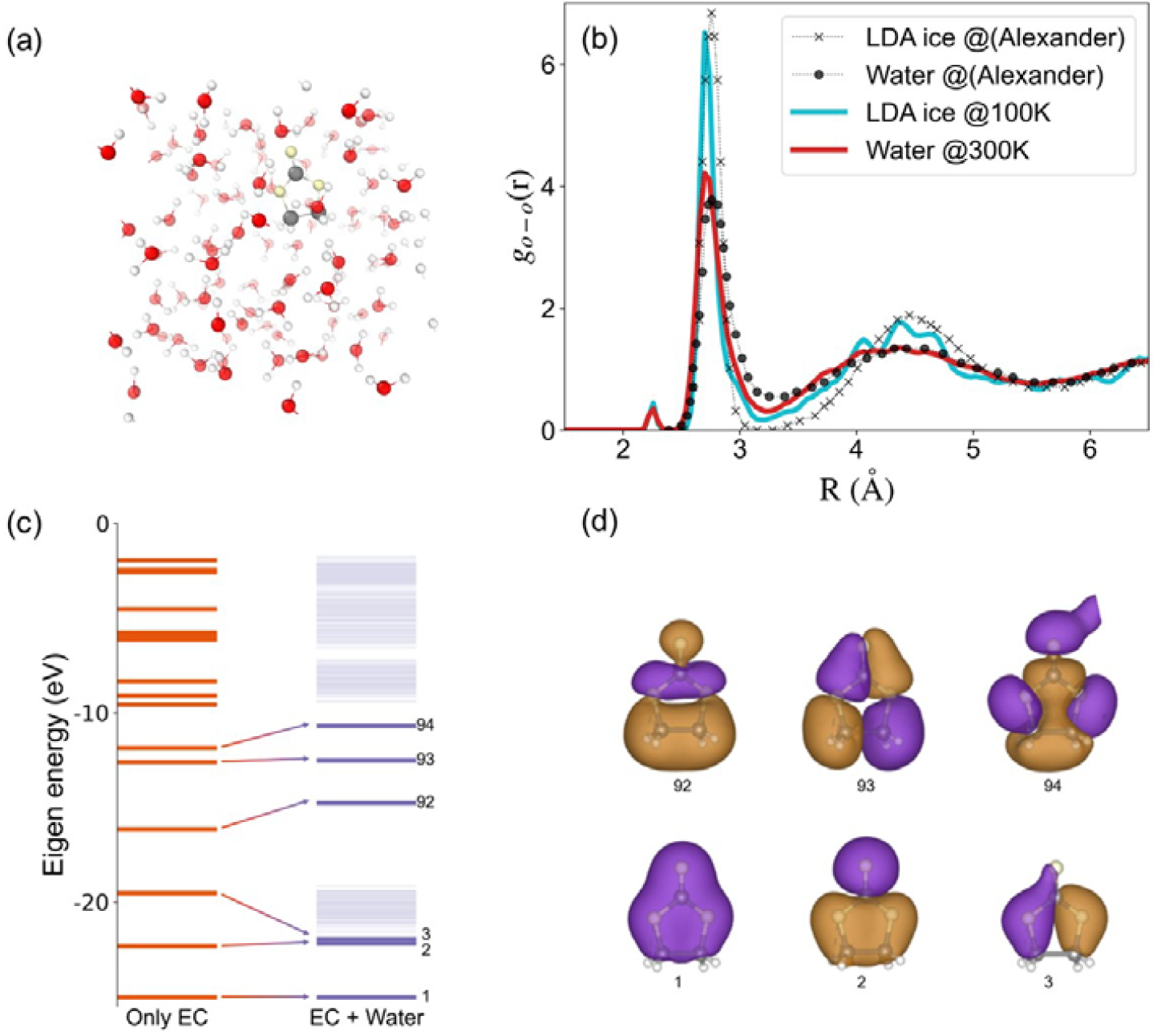
(a) The structure of the simulated sample under freezing conditions; the red and white parts indicate water molecules, and the brown and yellow parts indicate EC molecules. (b) Comparison of RDFs between oxygen atoms and oxygen atoms. The light blue and red solid lines denote our results, while the stars and dots are the results of Alexander *et al*.^50^. (c) Density-functional eigenvalue energy spectra of neutrally charged EC molecules and the simulation system. (d) Wavefunctions of some of the lowest-energy eigenstates of the EC molecule alone (numbered 1, 2, 3, 92, 93, and 94), while those higher-energy states (P-orbital electrons of the carbon and oxygen atoms) are prone to mixing with water.

When a material is irradiated with an electron beam, the initial ionization process may be single-electron ionization or it may lead to more complex processes such as secondary ionization, exciton excitations together with ionizations, double or more electron ionizations, etc. However, for small molecules, the multipole excitation happens within one molecule might still be relatively small, so we intend in this paper to consider first the simplest case of single electron ionization. We conducted 21 NOB simulations at each temperature with the initial state of ionization of one electron of State-1, and both simulations were run with 200 fs rt-TDDFT-MD duration with identical initial ionization conditions. After 200 fs, the hole might or might not have reached the HOMO (State-369 in Fig. 2) of the system; nevertheless, the reaction and wavefunction collapse (WFC) processes became rather slow. The purpose of this study is to determine whether differences in temperature (e.g., due to differences in Boltzmann distributions and detailed balances and due to differences in water environments) can lead to differences in initial molecule breakdown. Note that in theory, the temperature can lead to different reaction paths (both orbital coupling and WFC) and whether a WFC is allowed due to the energy conservation requirement. The stochastic branching nature of the simulations led to varying fragmentation results in the 21 NOB simulations at each temperature. In Fig. 2, we show the results of 4 NOB simulations, where (a)-(d) are under frozen conditions and (e)-(h) are at room temperature. The variations in the bond length, energy level and hole occupation observed in these four simulations are depicted. In Fig. 2 (a), (c), (e), and (g), three trajectories with different bond breaking patterns are shown, which are all from identical initial conditions except for temperature, demonstrating the stochastic nature of the simulations. In Fig. 2 (b), (d), (f), and (h), the corresponding energy levels and hole occupancy are also shown. At the beginning of the simulation, in all four cases, valence state-1 is occupied by one hole. Then, as the hole jumps to different energy levels, the bond lengths begin to change. The detailed bond breaking process is discussed below. Our simulations show that bond breaking is always caused by WFC events, typically between two deep levels with large energy separations. During one WFC, we observed both one bond breaking and multiple bond breaking. Notably, the bond breakings of some particular bonds are typically correlated with the hole transition between two particular states. It is easy to see that there are three bond lengths that increase significantly (indicating the breaking of such bonds): *C*_1_ - *O*_2_, *C*_1_ - *O*_3_ and *C*_2_ - *C*_3_, as shown in Fig. 2; only the time of the breaks and the mode of the breaks differ. From the point of view of the hole movements, there is no significant change in the bond lengths when the hole transits from State-1 to State-2 or State-3. It is not until the hole jumps to State-92 that the bond length begins to change dramatically.

**Fig. 2.**
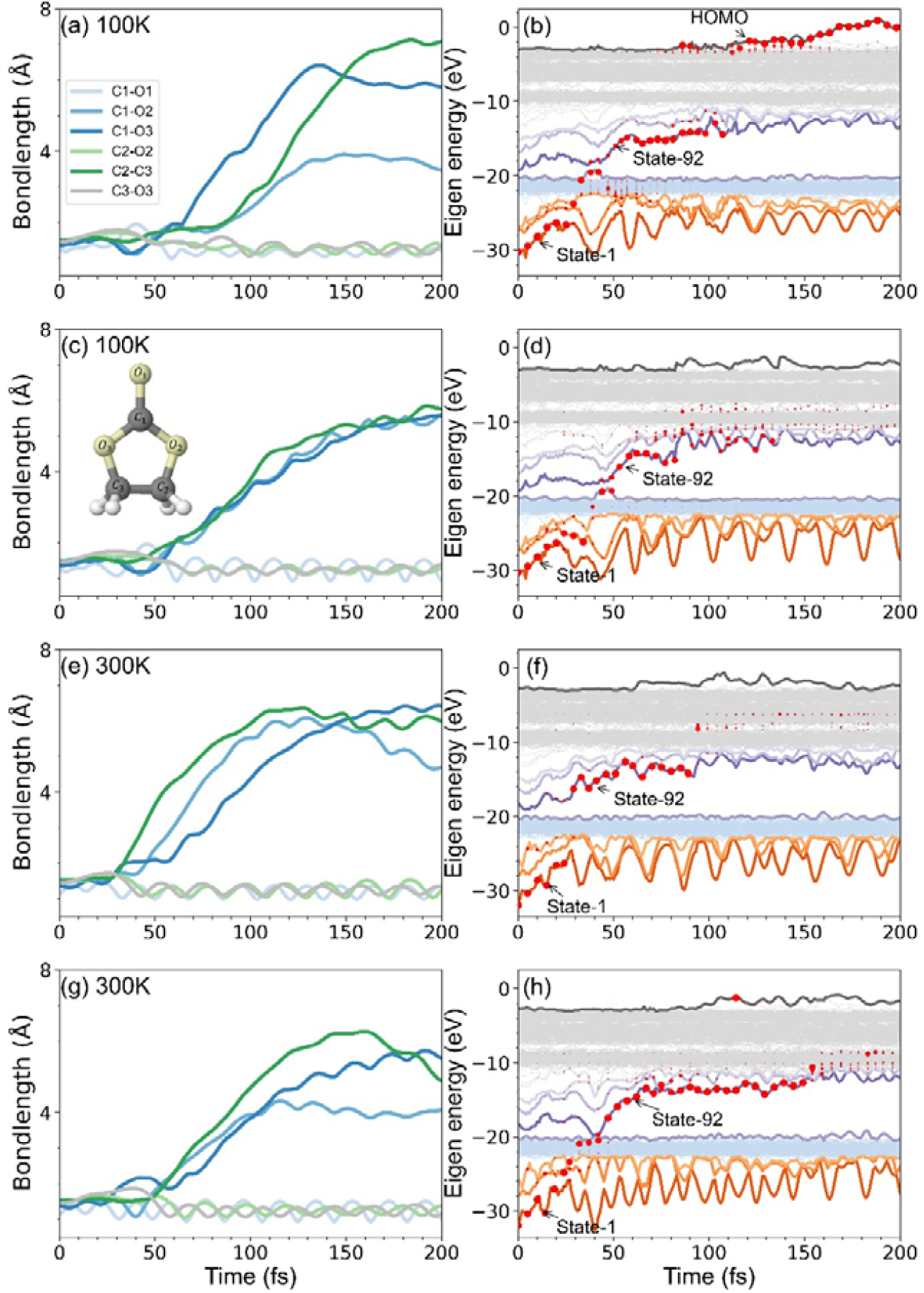
The left figures (a, c, e, g) display the changes in bond lengths in the EC molecule. Random numbers differ between (a) and (c) at 100K, and between (e) and (g) at 300K, resulting in different collapses as described in the ‘NOB method’ section. The EC molecular model and its atomic numbering are shown in (c). On the right, the figures (b, d, f, h) indicate the changes over time in the adiabatic state electron occupation and energy levels of the entire system. The lines indicate the energy level changes, and the red dots represent the occupation of holes, with the size of the dots being proportional to the number of the hole occupied.

We found that there may be a relationship between hole transition and bond breaking. Fig. 3(a) illustrates the conclusions we obtained by summarizing the results of all 42 simulations. We divided the energy levels of the system into four parts. The first part included State-1, State-2 and State-3; the second part included water molecules, which represented deeper energy levels (State-5 to State-90); and the third part represented State-91, State-92, State-93 and State-94 because we found that with the transition of the hole, it caused State-91 to become an eigenstate of the EC as well. The fourth part represents the shallower energy levels, which are the mixing of EC and water molecules (State-95 to State-369). We found that the hole jumps from the first part to the second part with 14.3% and 12% probabilities under freezing and room temperature conditions, respectively, and this process hardly leads to bond breaking. Moreover, the transition to the fourth part, with probabilities of 14.3% and 28%, leads to a large change in the structure due to the large energy difference, but it is uncertain which bonds will change. The probabilities of transfer to the third part are 71.4% and 60%, respectively, and we found that when the hole jumps to State-93 and State-94, breakage of *C*_1_ - *0*_2_ and *C*_3_ - *O*_3_ occurs under two temperature conditions. When the hole transitions to State-91, it results in the breakdown of three types of bonds, namely, *C*_1_ - *O*_1_, *C*_2_ - *O*_2_ and *C*_3_ - *O*_3_. The transfer to State-92 almost causes the breakage of *C*_1_ - *O*_2_, *C*_1_ - *O*_3_ and *C*_2_ - *C*_3_, as shown in Fig. S2-4 in the Supplemental Material. We also have performed several additional simulations of ionizing state-2 and state-3 and found similar patterns, with some connection between electron jumps and bond breaking as shown in Fig. S5 and Fig.S6 in the Supplemental Material. We therefore believe that there may be some connection between the state transition and bond breaking. Finally, we also summarized the final fragment information, as shown in Fig. 3(b). The blue and red lines in the figure mainly indicate the probabilities of the most common fragments (*O, O* - *C* - *O, C* - *C, C* - *O, C* - *C* - *O*), with “Other” indicating the less common fragments. We found that temperature has only a small effect on the initial stage of molecule breakdown and fragment generation. The two temperature systems have very similar molecule breakdown patterns and magnitudes.

**Fig. 3.**
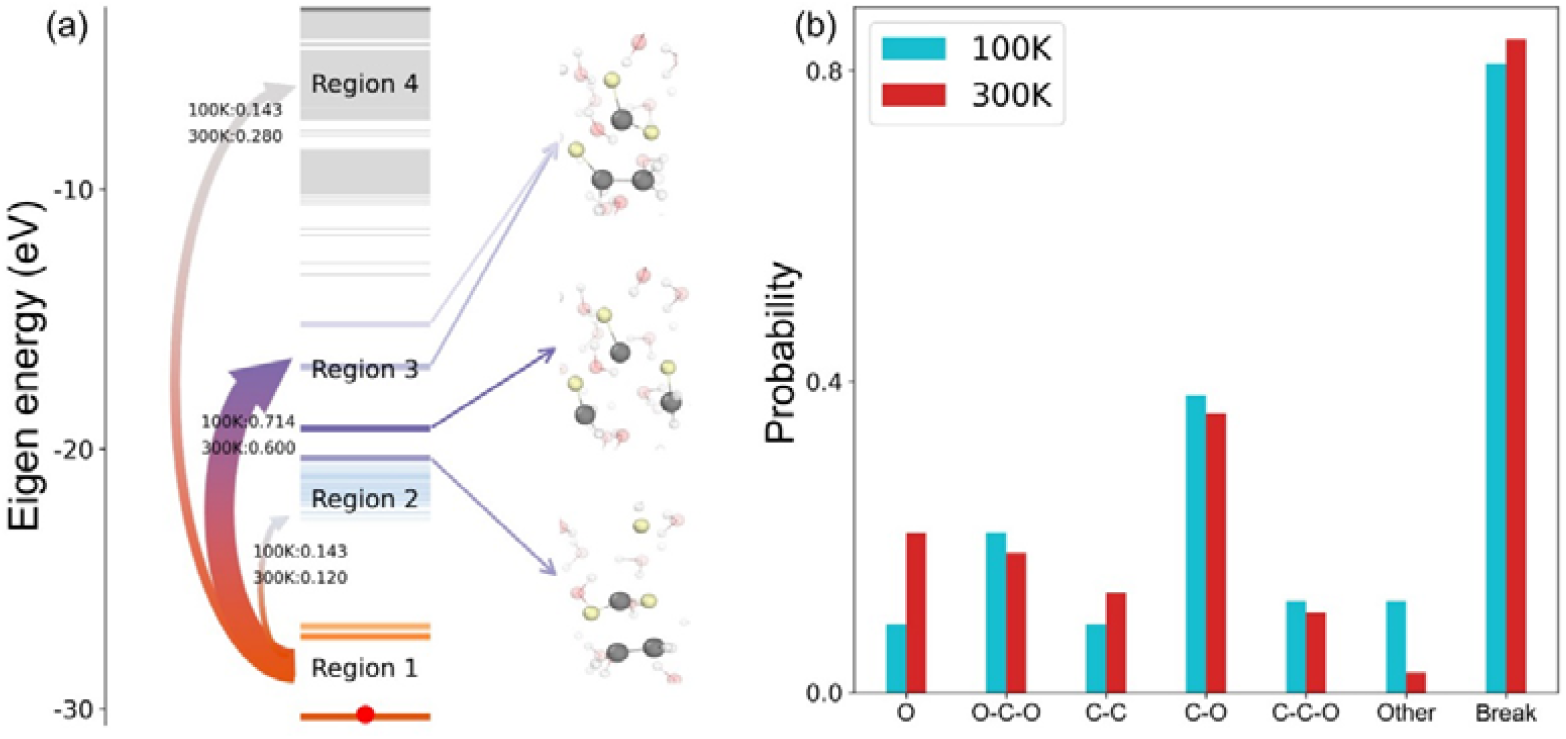
(a) The path of hole transitions and the pattern of molecule breakdown in all the simulations; the red dots represent the occupation of holes. The left panel shows the probability of the hole jumping from Region 1 to the other Region at two temperatures. The right image shows the resulting molecular fragmentation because the hole jumps to Region 3. (b) The number of fragments and the percentage of bond breakage that occurred. “Other” refers to the case of less common fragments. The “Break” columns indicate the probability of at least one bond breaking during the 200-fs simulation.

The above results illustrate that the impact of electron beam irradiation on molecular dissociation in the physical stage is consistent at both temperatures. What happens next is crucial. Note that during the above NOB simulation, the hole transition energy is transferred to the fragments. Thus, the typical fragments will have a large kinetic energy. We will see what happens for these fragments at these two temperatures. We carried out 2-picosecond ab initio molecular dynamics (AIMD) simulations following each of the 200 fs TDDFT-MD simulations (using the final atomic positions and velocity of the TDDFT-MD as the initial conditions for the AIMD simulations). For the AIMD, we used a neutral system, assuming that the ejected electron returned to the molecule. We first found that at both temperatures, there are significant second-stage reactions (secondary reactions) between the fragments and water molecules. In summary, in high-temperature cases, most secondary reactions occur between the fragment and surrounding water molecules. The fragments have sufficient kinetic energy to break into the surrounding liquid water and to react with the liquid water molecules. On the other hand, at low temperature, the surrounding water molecules freeze, and the fragments bounce back; thus, there is a cage effect^9,12,53^. As a result, the reactions occur only amount the fragments; thus, the fragments tend to repair and go back to the original molecule. The results for 100K and 300K are shown in Fig. S8 in Supplemental Material, and we additionally performed 4K simulations, the results of which are shown in Fig. S9 in Supplemental Material. Let us first use the evolution of the “O” radical presented at the end of the TDDFT simulation to illustrate this point. At 300 K, the “O” fragments move away from the original EC location, easily combine with water molecules to form hydrates, and even adsorb the H atoms of the water molecules to themselves. On the other hand, at 100 K, the “O” fragments remained close to the EC location and close to the other fragments. It reacts with other fragments to form new fragments. This situation is illustrated in Fig. 4 (a) and (b). At the end of the AIMD simulations, for both temperatures, all the O fragments react with other species, but their reaction patterns are very different.

**Fig. 4.**
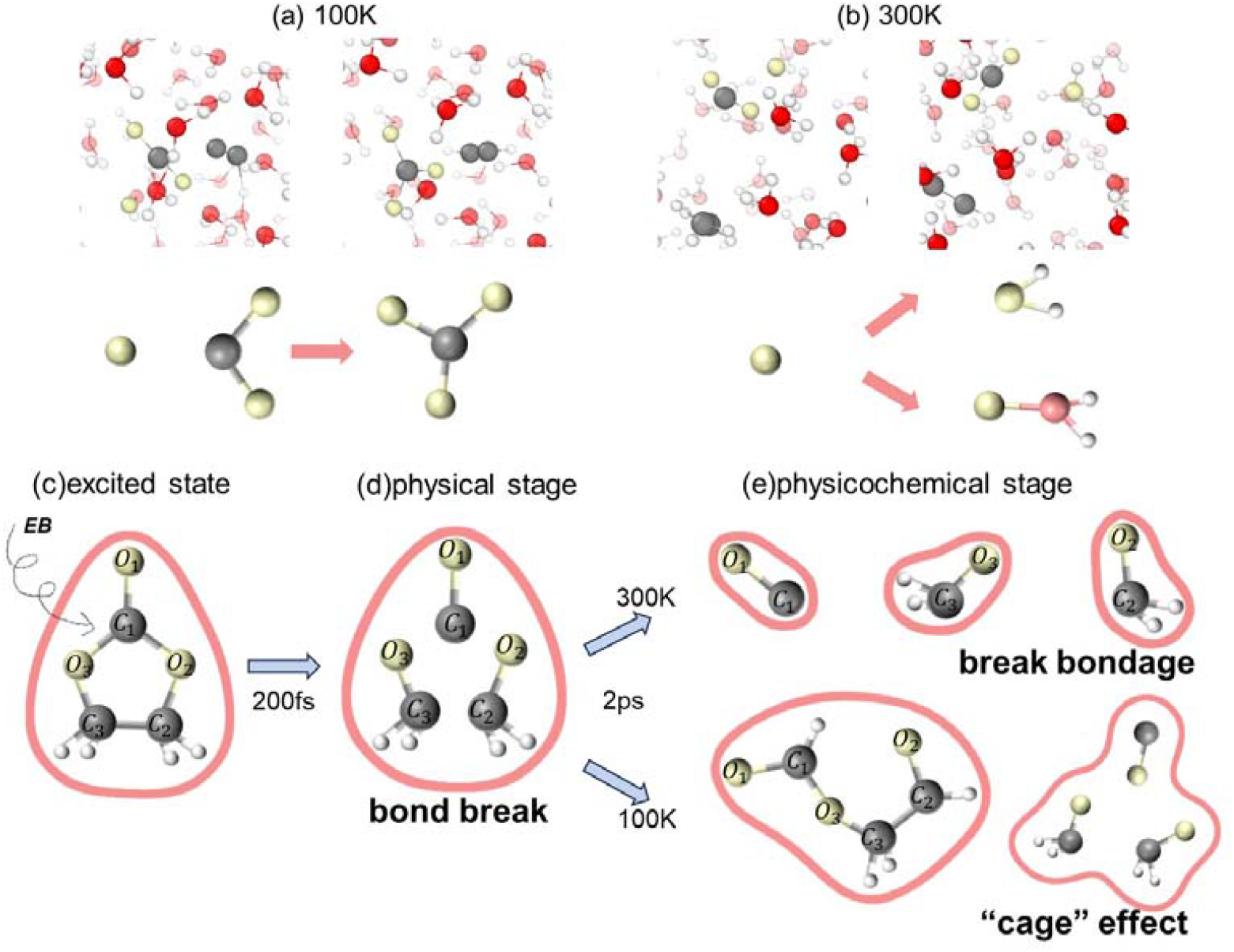
Dynamical processes of fragments. (a)(b) Paths of individual oxygen atom fragments in AIMD simulations. Case (a) occurs in the 100 K, while (b) occurs in the 300 K. (c)(d)(e) Dynamics of EC molecules after ionization for 2 picoseconds under freezing and room temperature conditions. The excited state (c) leads to bond breaking, which is reflected in the physical stage (d); finally, the physicochemical stage (e) shows the difference between the two conditions, where the fragments are reorganized during freezing due to the “cage” effect and break the cage at room temperature. The yellow atoms indicate oxygen atoms in EC molecules, the brown atoms indicate C atoms in EC, the red atoms represent oxygen atoms in water molecules, and the white atoms represent hydrogen atoms in water molecules. Abbreviations: electron beam, EB.

The cage effect also occurs for other fragments. For example, at 100 K, fragments “*C*_2_ - *O*_2_” and “*O*_l_ - *C*_l_ - *O*_3_ - *C*_l_” will combine to form a long chain, which has a curved shape similar to the original EC, as shown in Fig. 4 (c), (d), and (e). In our 2-ps AIMD following the rt-TDDFT simulations, we did not see the long chain fully returning to the EC. Thus, the healing of the damage is only partial. To estimate the possibility for the chain to fully return to the EC, we used the NEB method to calculate the reaction barrier needed to form the EC (the EC energy is approximately 1 eV lower than that of the chain). For this to occur, the C1 and O2 atoms must move closer together to form the C1-O2 bond, and at the same time, one hydrogen atom must jump from the C1 atom to the C2 atom. This has a relatively large barrier of 1.5 eV. Thus, it might take a long time for this to occur. We find that at the end of our 2-ps AIMD, the molecule has a temperature of 800 K. If this temperature remains constant, the reaction back to the EC can occur at approximately 0.003 seconds. Overall, by combining all these simulation results, we can conclude that the reduced damage caused by low-temperature TEM is due to the cage effect.

## Conclusion

In conclusion, we investigated the molecular dissociation process and fragmentation kinetics of small organic molecules irradiated by electron beam using RT-TDDFT and AIMD. Images of fragmentation generation after ionization of EC molecules and subsequent fragmentation reorganization under different temperature conditions were obtained, as shown in Fig. 4. We found that the dissociation processes of molecules immediately after initial ionization are almost identical, with similar fragmentation and final product probabilities. The different nuclear vibrations in this period do not play any significant role. Perhaps one of the reasons is that the molecule is quickly heated by the hot carrier cooling energy; thus, the initial temperature does not play any role. The movement of the surrounding water molecule also does not play a significant role, probably because during the dissociation process, the hole mostly remains in the molecule. However, after the initial dissociation process, in the second longer time scale period, where the molecular fragments are in electronic adiabatic ground states, the frozen water molecules in the low-temperature case play an important role. They can prevent fragments from flying apart and reacting with water molecules (as in the high-temperature case). This cage effect keeps the fragments together, thus facilitating their back-reaction. We did observe some partial repair reactions in our follow-up AIMD simulations; however, to fully return to the EC, a much longer time is needed than our AIMD simulation can afford. Overall, in Cryo-TEM, this is not because there is no initial damage but that secondary chemical reactions and the fate of fragments produced by radiolysis are constrained by the cage effect.

## Methods

### The simulation settings

In this study, we performed real-time simulations of time-dependent density functional theory molecular dynamic simulations(rt-TDDFT-MD) and ab initio molecular dynamics (AIMD) simulations using Perdew–Burke–Ernzerhof (PBE) exchange correlation functions^54^. The computations were executed with spin polarization and relied on the PWmat code^48,49^, which is implemented on graphics processing units (GPUs) using plane-wave pseudopotential Hamiltonian. We use an SG15 optimized norm-conserving Vanderbilt pseudopotential (ONCVPSP), and the basis set of plan wavefunctions is obtained using an energy cut-off of 50 Ry. The same initial ionization process at both temperatures can be calculated using the binary encounter-dipole (BED) formula^19^. A sudden transition approximation is taken, which means that the atomic configurations (and their velocities) are still in a neutral state, and Kohn–Sham electron wavefunctions {*ϕ*_i_} are still in the neutral charge state at *t* = 0. Note that after removing one electron in *ϕ*_*i*_, the remaining {*ϕ*_*i*_} are no longer actual eigenstates. We then carry out the NOB calculations, and our adiabatic state-based algorithm^52,55^ allows us to use a relatively large time interval Δt (0.1 fs).

### NOB method

In most TDDFT calculations, the Schrödinger equation is solved for by solving for the single-electron wavefunction. In the NOB, the wavefunction evolution is described by the Liouville equation of the density matrix *d*(*i,i*′,*t*), which is constructed on a basis set of single-particle adiabatic eigenstates {*ϕ*_i_ (*t*)}:

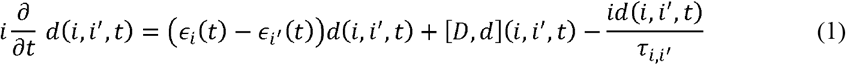

Here, ∈_*i*_(*t*) is the eigenenergy of the adiabatic state *ϕ*_*i*_(*t*), the communicator [*D, d*] is defined as [*D, d*] = *Dd* − *dD*, and the adiabatic coupling constant is defined as:

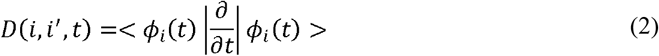

Note that in Eq. (1), the decoherence effect is described by the decay of the off-diagonal element *d*(*i,i*′,*t*) with respect to the dephasing time *τ*_*i,i*′_. For diagonal elements *i* = *i*’, *τ*_*i,i*′_ is infinite. In theory, *τ*_*i,i*′_ can be calculated from the dynamics itself^56^. It can be shown that, under Eq. (1) and the usual Heyman–Feynman force calculation, the total energy of the dynamics is conserved. On the other hand, the density matrix can be diagonalized, with its eigen vector defined as the natural orbital and eigenvalue being *λ*_*k*_(*t*) ∈ [0,1]:

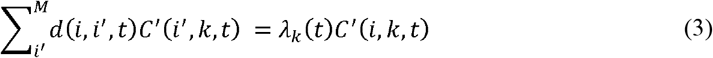

The natural orbitals are defined as follows:

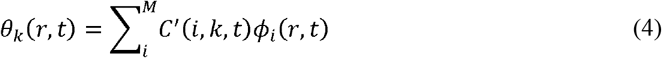

Due to the decay of the off-diagonal term of the density matrix, *λ*_*k*_ (*t*) is no longer 0 or 1, representing the ensemble nature of the density matrix. One can use an entropy expression to measure the degree of such an ensemble nature:

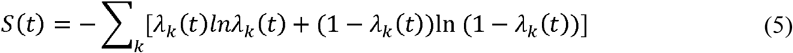

If *s*(*t*) is not 0, then the density matrix does not consist of a pure quantum mechanical state (a single Slater determinant (SD)); instead, it is a sum of several decoherent Slater determinants (SDs). In the NOB scheme, if *S*(*t*) becomes bigger than a critical value (*S*_*c*_), we will stochastically collapse the system into a single Slater determinant consisting of a set of natural orbitals *θ*_*k*_ (*r,t*), as shown in Fig. S6 in the Supplemental Material. In practice, after each integration step Δ*t*, the value of *S*(*t*) is computed; if *S*(*t*) > *S*_*c*_, a wavefunction collapse (WFC) event will occur, a set of N *θ*_*k*_ (*r,t*) will be occupied, and the dynamics will be restarted from this new state with *S*(*t*) =0. These N natural orbitals *θ*_*k*_ (*r,t*) are chosen stochastically with a probability proportional to *λ*_*k*_ (*t*) After the WFC, these chosen natural orbitals will have an occupation 1, while all the other natural orbitals will have the occupation 0. Additionally, the change in energy of the electronic structure will be transferred to nuclear kinetic energy on the transition degree of freedom. If the transition results in an increase in the energy of the electronic structure, then the nuclear kinetic energy of this transition degree of freedom (TDF) will be reduced to compensate for the increased energy. If the kinetic energy in this degree of freedom is insufficient, then this attempted transition will be abandoned. This approach will restore the detailed balance.

The transition degree of freedom (TDF) and its corresponding force (impact on the nuclear momentum) for one WFC is defined as follows:

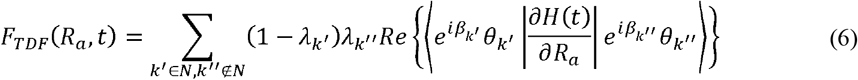

Here, *β*_*k*′_ and *β*_*k*″_ are two stochastic phases chosen for states *k*′ and *k*″.A transition impulse is provided to the momentum along this degree of freedom (DOF) to satisfy the total energy conservation: *V*’(*R*_*a*_,*t*) = *V* (*R*_*a*_,*t*) + *αF*_*TDF*_ (*R*_*a*_,*t*) /*M*_a_, where *M*_a_ is the mass of atom a. If we define the change in the electronic structure energy after the WFC as 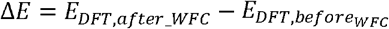, then if Δ*E* < 0, the WFC is always accepted. If Δ*E* > 0 and there is no solution for α to satisfy the total energy conservation (there is not sufficient kinetic energy in this transition degree of freedom), then this WFC will be rejected. As described above, this approach guarantees a detailed balance, making the cooling process much more favorable than the heating process.

## Data availability

The data that support the findings of this study are available from the corresponding author upon request. Source data are provided with this paper.

## Acknowledgements

We thank Dr. Xiao-Xiang Yu and Bo Chen for helpful discussions. This work was supported by the National Natural Science Foundation of China under Grant Nos. T2293700 and T2293702 and the Science and Technology Innovation Program of Hunan Province under Grant No. 2021RC4026.

## Author Contributions

L.W.W. designed the project. J.Y.D. and L.W.W. suggested the specific scientific problem and the general idea on methodology. Y.L. performed the TDDFT-MD simulations and analysed the data. Y.L., D.D.K., J.Y.D., and L.W.W. interpreted the results. Y.L. wrote the paper, and D.D.K., J.Y.D., and L.W.W. edited the manuscript before submission. All the authors critically evaluated the manuscript.

