## Supplemental Material for "The cage effect of electron beam irradiation damage in cryo-electron microscopy"

**Supplemental Material: The cage effect of electron beam irradiation damage in cryo-electron microscope**

**Structural simulation process**

We first performed NVT relaxation of the simulated structure at 1000 K for 1000 fs. Then the relaxed structure was used as the initial structure and annealed to 100 K and 300 K, respectively, with an annealing time of 10,000 fs. Finally, the annealed structures were used to perform the NVT relaxation process up to 10,000 fs at 100 K and 300 K, respectively. The time step of all processes is 1 fs.

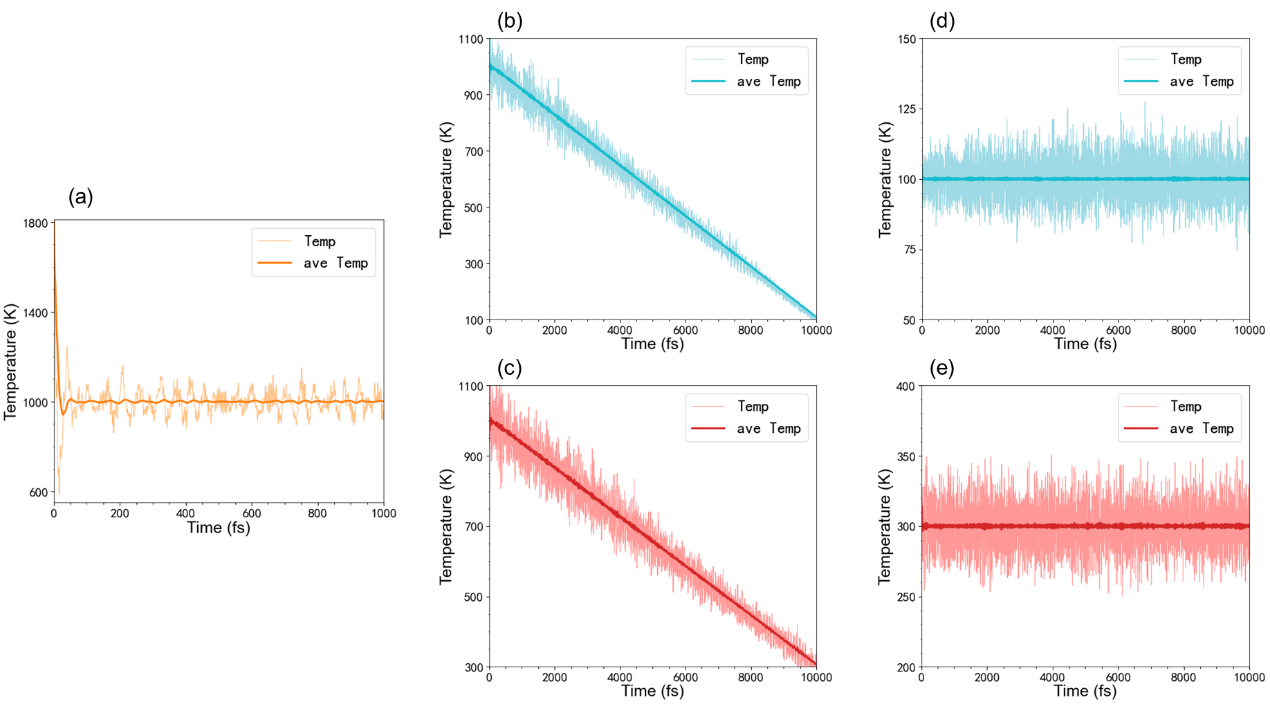

Fig. S1 Temperature variation during structural simulation. Temperature changes during relaxation at 1000 K(a), 100 K(d) and 300 K(e). Temperature changes during annealing at 100 K(b) and 300 K(c).

**NOB simulation results**

The Fig. S2, Fig. S3 and Fig. S4 show the bond breaking that occurs when the hole moves. It is important to note that in the diagram, $C_{2}-O_{2}$ and $C_{3}-O_{3}$ are equivalent due to the left-right symmetry of EC's structure. Similarly, $C_{1}-O_{2}$ and $C_{1}-O_{3}$ are also equivalent.

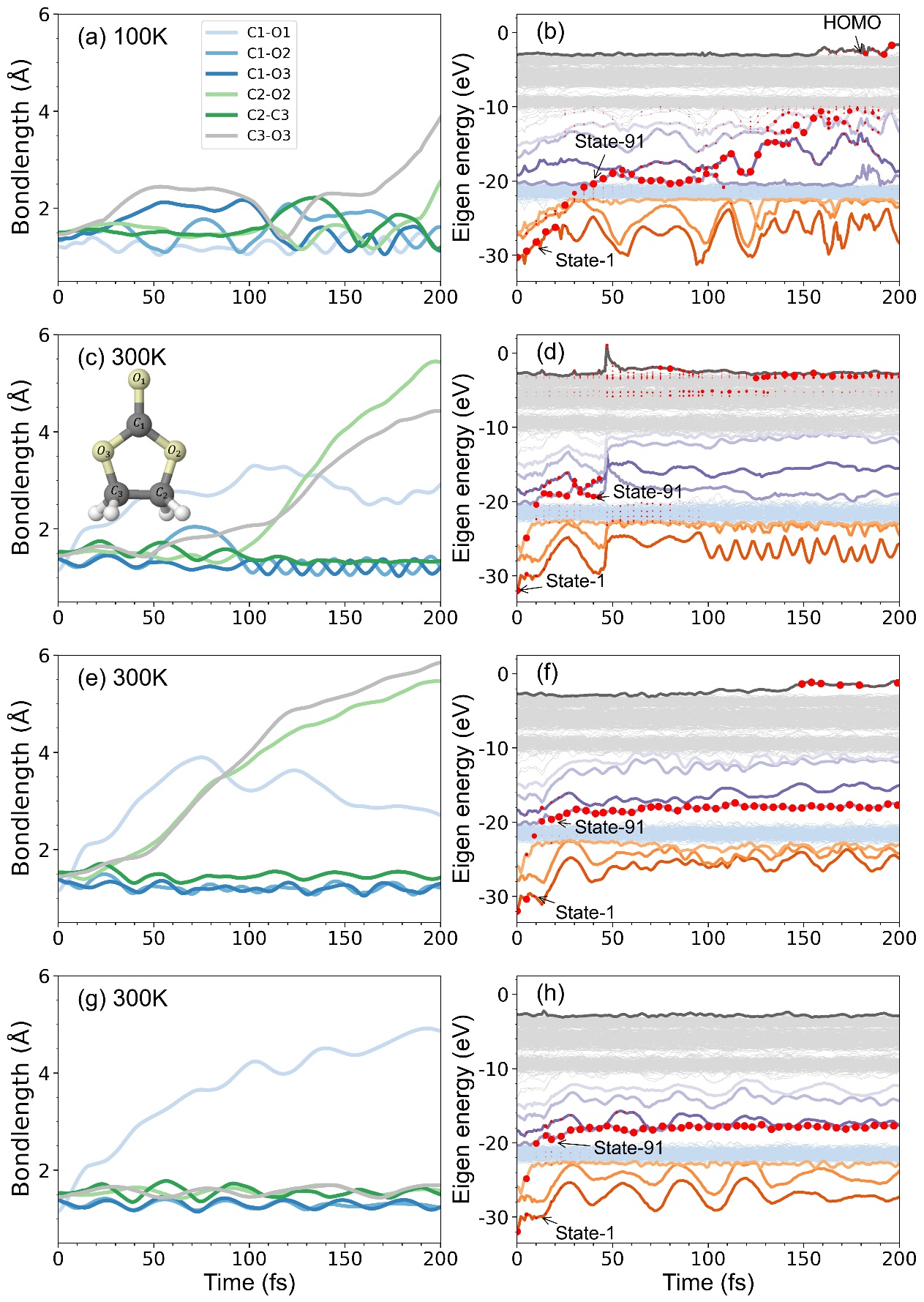

Fig. S2 The bond length changes corresponding to the hole jump to State-91 and the energy level change.

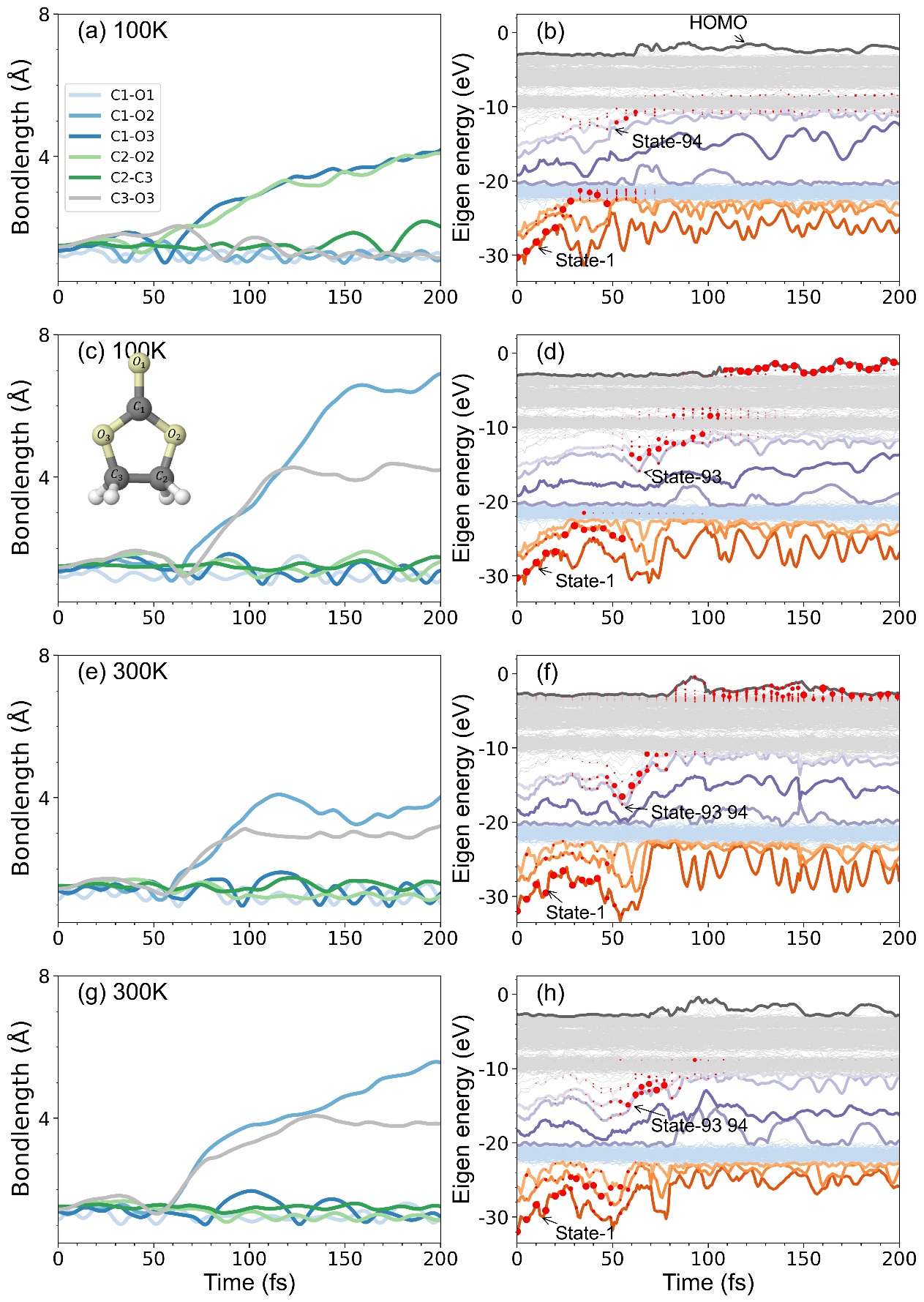

Fig. S3 The bond length changes corresponding to the hole jump to State-93 and State-94, and the energy level change.

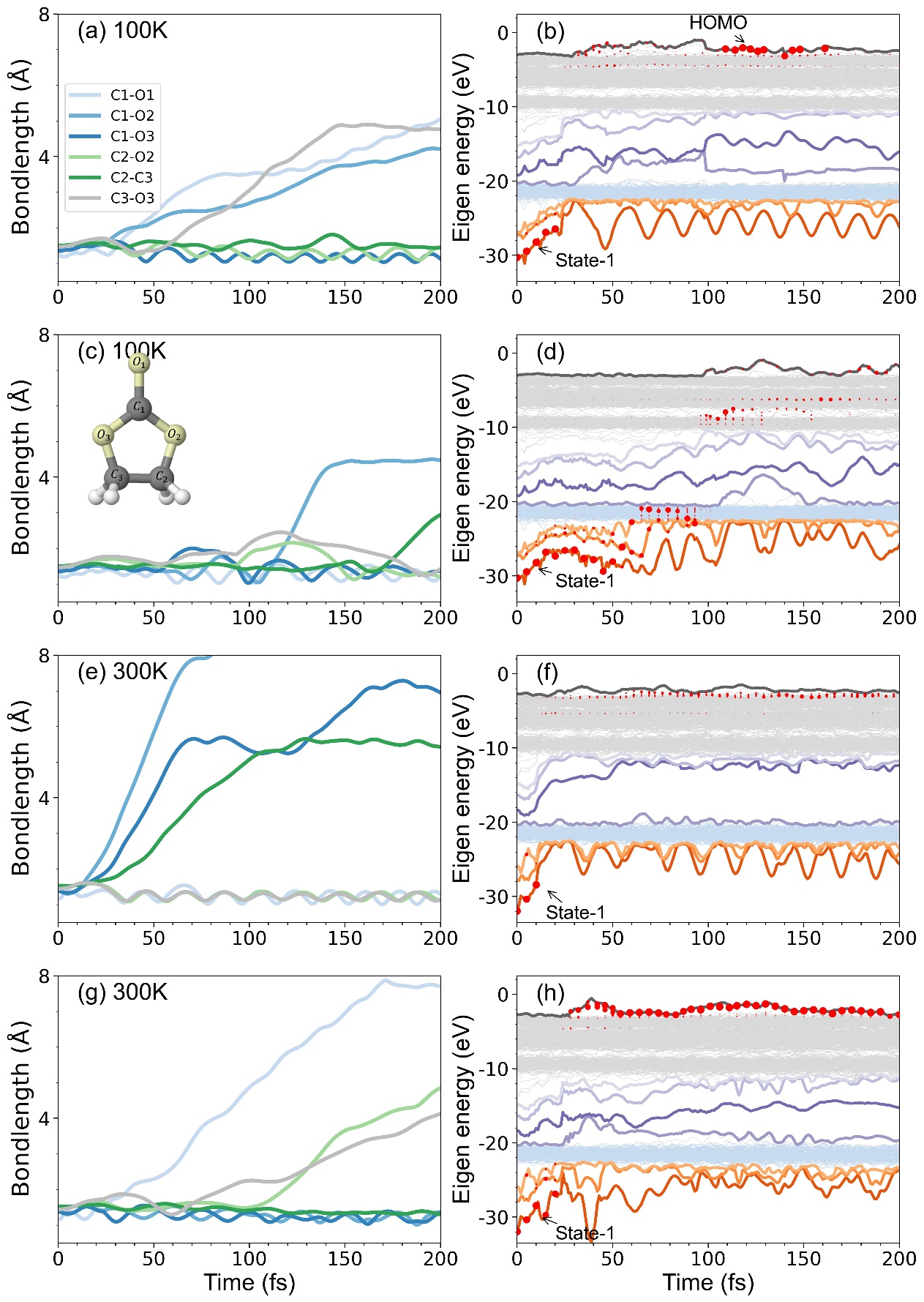

Fig. S4 The bond breaking that occurs when the hole moves to State except State-91, 92,93, 94.

The Fig. S5 and Fig. S6 show bond length changes and hole-leaping scenarios when the initial ionization is State-2 and State-3, respectively.

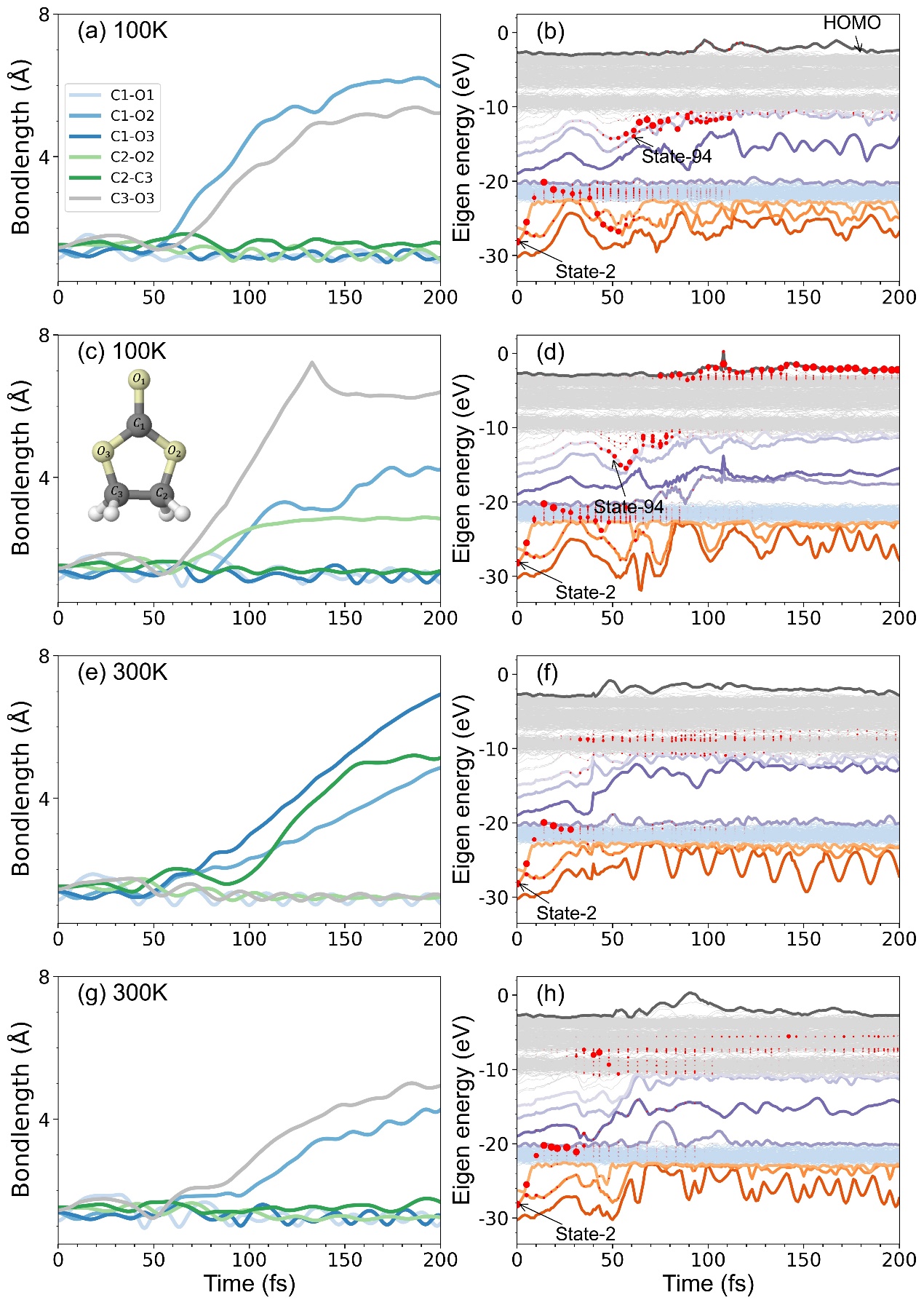

Fig. S5 Changes in the bond length of the system after ionization of state-2 energy level electrons and changes in holes.

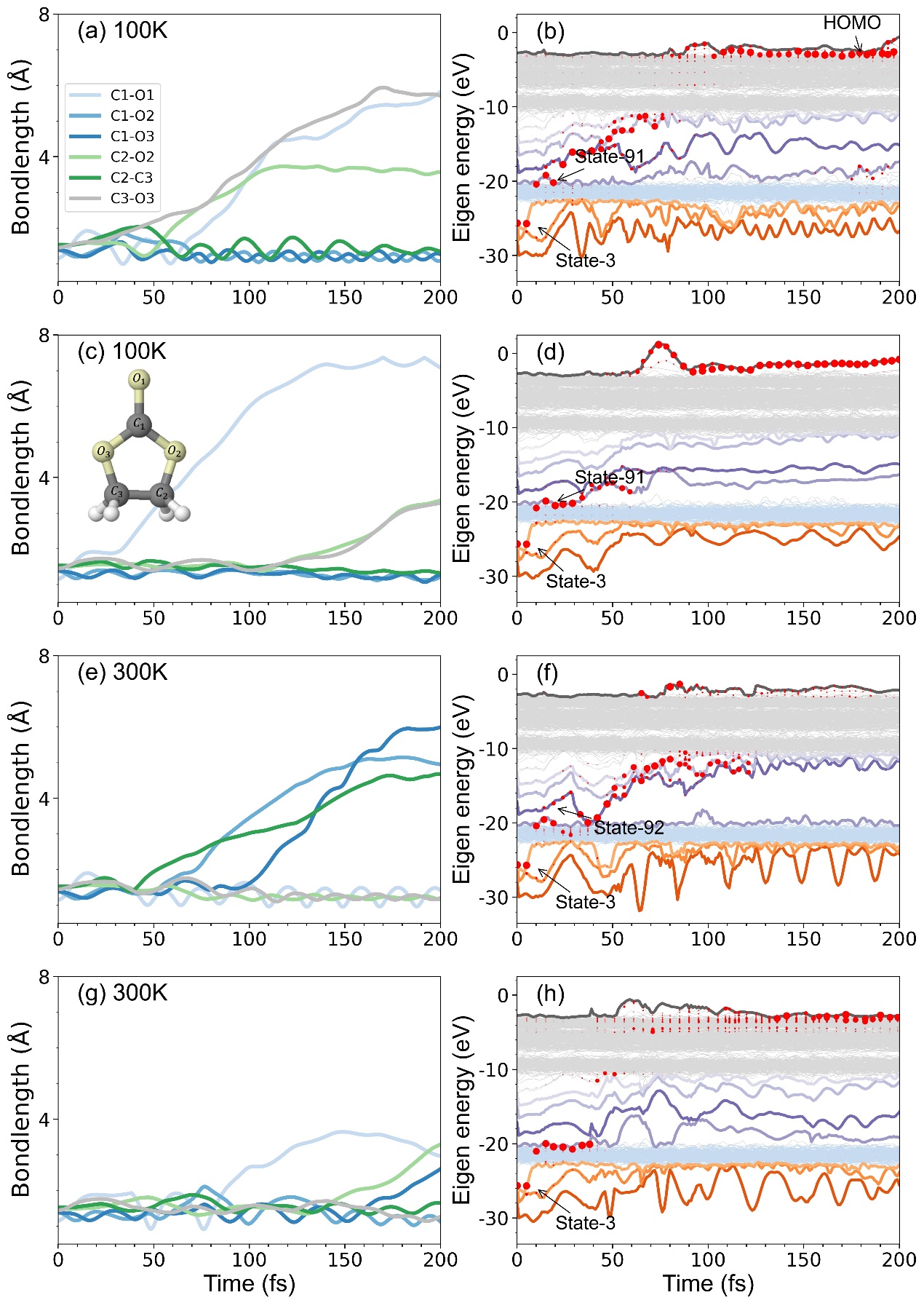

Fig. S6 Changes in the bond length of the system after ionization of state-3 energy level electrons and changes in holes.

Fig. S7 illustrates the variation of the system for different simulation sizes.

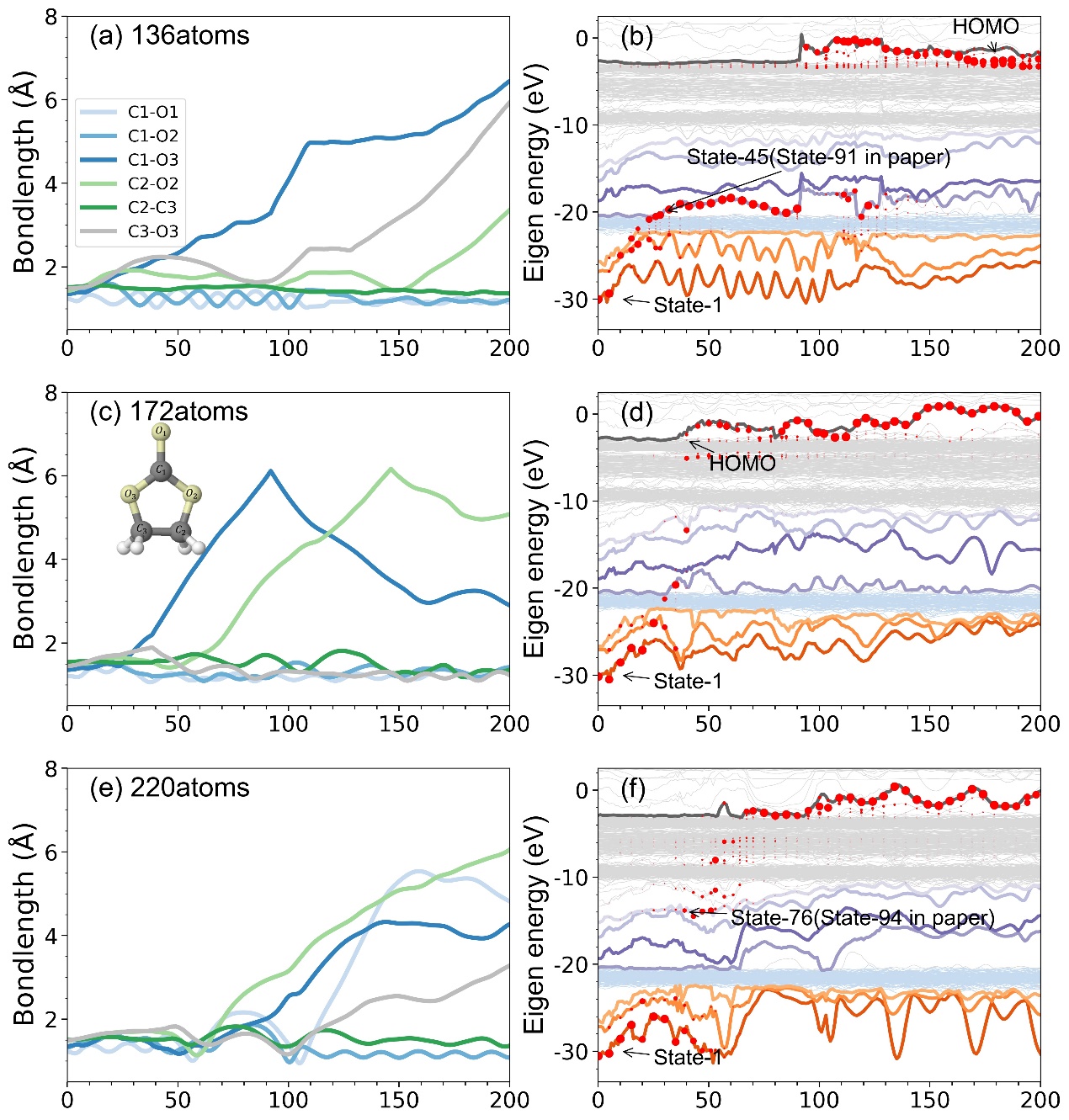

Fig. S7 Bond length variations and hole-leaping processes in the simulated regime with different dimensions. (a)(b) for 136 atoms, (c)(d) for 172 atoms and (e)(f) for 220 atoms.

**AIMD simulation results**

Some snapshots of the final structure of some of the AIMD simulations at 100 K and 300 K are shown below. From the figure, we can roughly see that at 100 K, the fragment molecules are more concentrated and there are no water molecules in between the fragments, but for 300 K, the distribution of fragments is more dispersed and there are water molecules in the middle of the fragments. At 300K, we also found a reorganization of fragments as shown in Fig. S8(i). But the fragments were mixed with a water molecule during the reorganization.

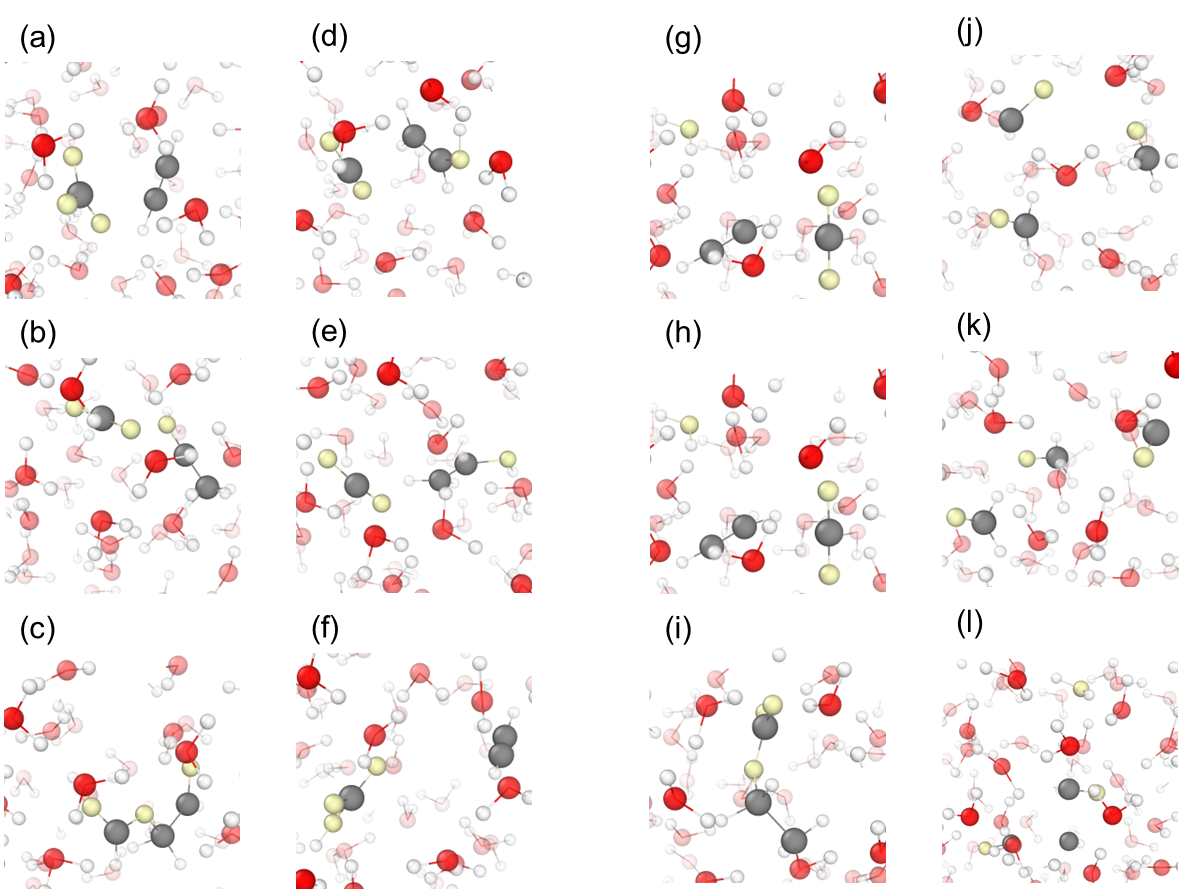

Fig. S8 AIMD final structural diagram. (a)(b)(c)(d)(e)(f) for 100 K, (g)(h)(i)(j)(k)(l) for 300 K.

Fig. S9 shows the AIMD results considering van der Waals effects, and we find that the fragments are more dispersed at 300 K, distributed almost throughout the cell space, while they are more concentrated at 100 K, moving near the initial position of the EC molecule. And we also performed 4K simulations and found that the fragments also moved near the initial position, but the motion would be slower compared to 100K, and there was no tendency for the debris to move closer to each other.

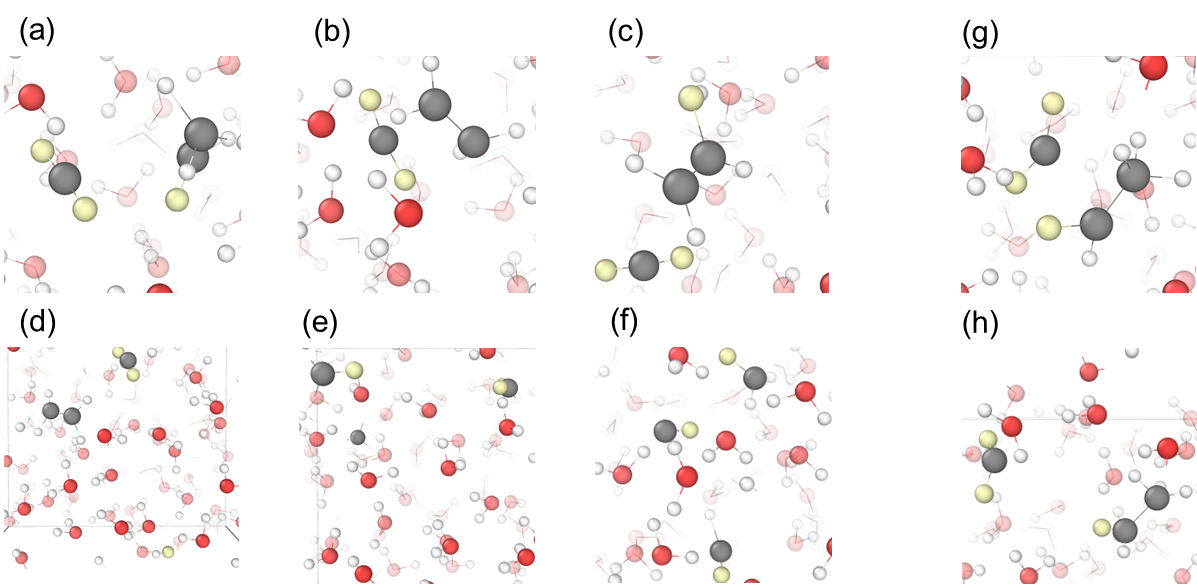

Fig. S9 AIMD final structure considering van der Waals effect. (a)(b)(c) for 100 K, (d)(e)(f) for 300 K. And (g)(h) for 4K.

**Natural orbital branching (NOB) Methods**

In Fig. S10 (a), we show the variation of entropy in the NOB with time when we set $S_{c}=0.5$ when at t=0 the system has a single hole in the lowest state of the valence band in one of its spin state, while we have used spin=2 LSDA calculations. Fig. S10 (b) demonstrates the variation of electron occupancy for each intrinsic adiabatic state in the simulation, it is simply defined as the diagonal matrix element $d\left( i,i^{'},t \right)$ of adiabatic state $\phi_{i}(t)$. We have set $\tau_{i,i^{'}}$ for all the off-diagonal part to be 10 fs. We found that the final result does not sensitively depend on this number. We find that at the initial 20 fs, there are fewer collapse events because the holes are mainly localized on state-1 at this time, and it is well separated in energy from other adiabatic states. After 20 fs, the hole jumps to higher energy levels, creating many partial holes in other adiabatic states. After this, a dense period of WFC appears, with the system quickly increase its $S\left( t \right)$ after each WFC. This happens until 100 fs, when the holes are stabilized in the HOMO energy level, and the number of collapse events begins to decrease significantly.

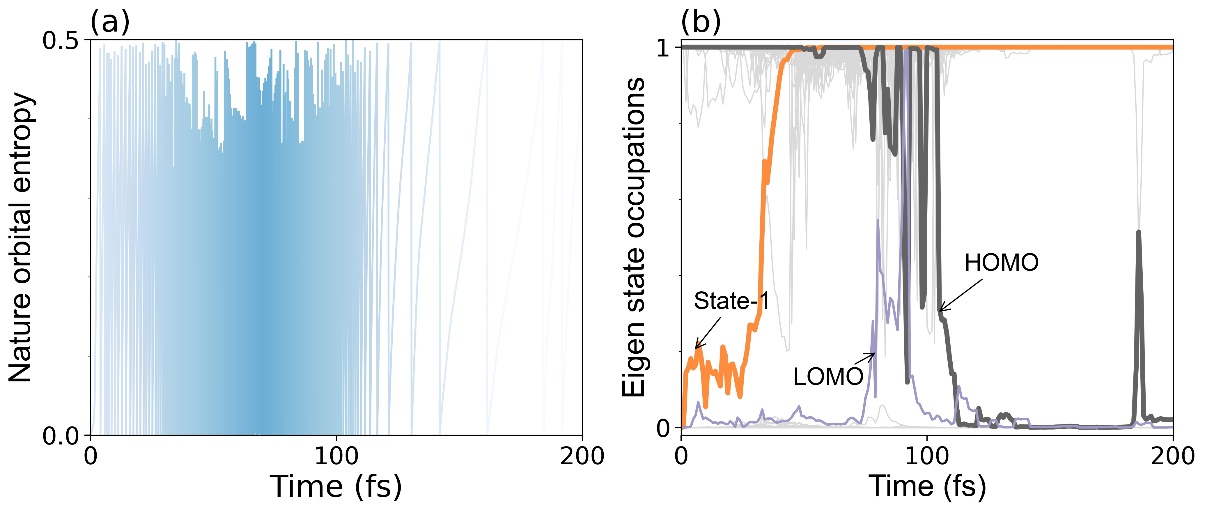

Fig. S10 Ionization of state-1 at t = 0 for $S_{c} = 0.5$ (a) indicates the change in entropy with simulation time, (b) demonstrates the electron occupation of the eigenstates.

**Natural orbital branching (NOB) example**

Below we show an example to illustrate the change of the wave function and the change of the velocity of the C and O atoms in the EC during electron jumps, which corresponds to Fig. 2 (c)(d) in the paper. We can see that after 20 fs, the hole is transferred from state-1 to state-2, at which time the atomic velocity does not change much, but the shapes of the wave functions corresponding to state-1 and state-2 change; while after 40 fs, the hole jumps to state-91, at which time the atomic velocities, especially $C_{1}$ and $O_{1}$, change considerably and lead to bond breaking, and at the same time, the wave functions corresponding to state-91 also change considerably.

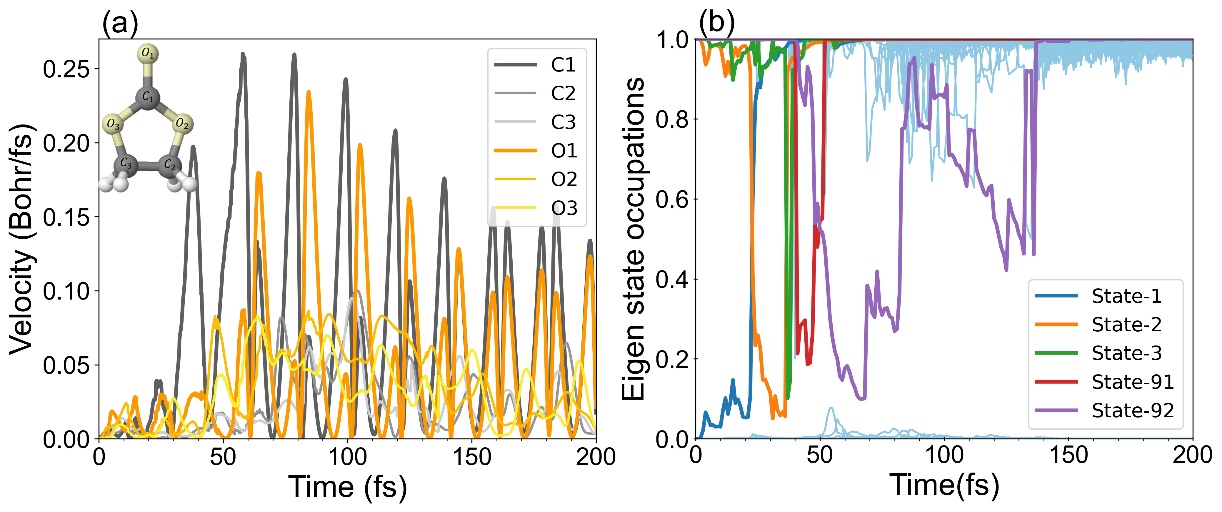

Fig. S11 The change in velocity of the C and O atoms in EC.

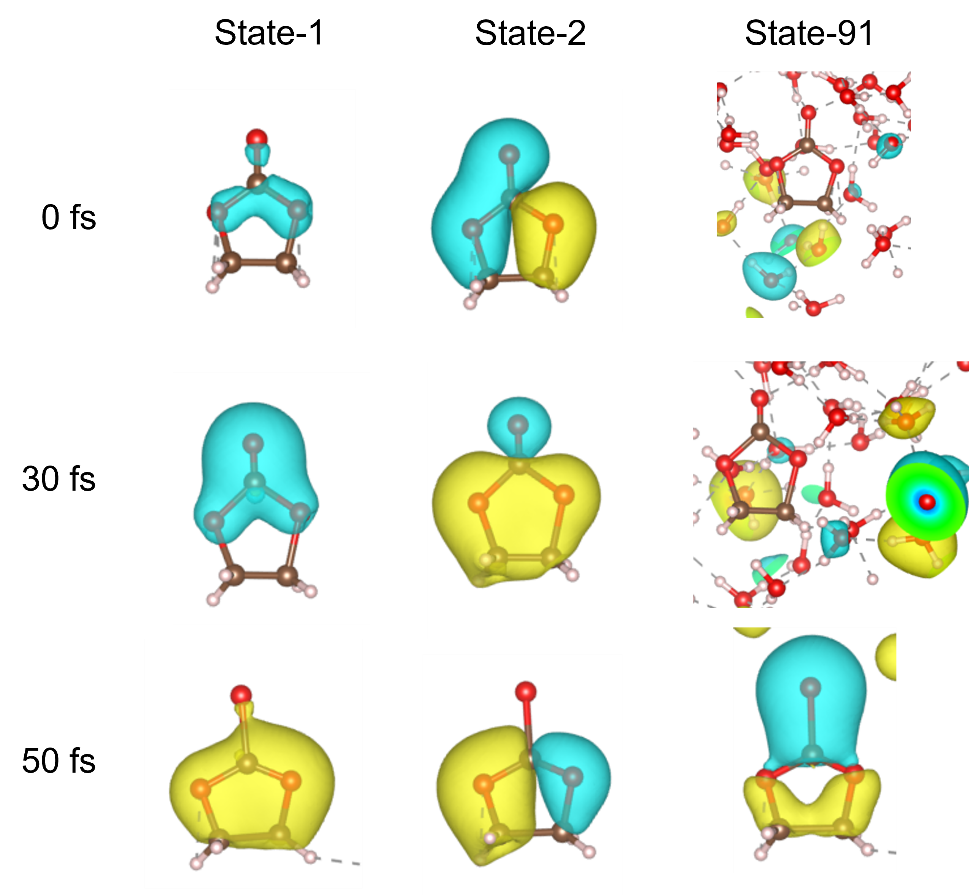

Fig. S12 The major change in the wave function after electron jumps.
